# Cloud-controlled microscopy enables remote project-based biology education in Latinx communities in the United States and Latin America

**DOI:** 10.1101/2022.08.05.502091

**Authors:** Pierre V. Baudin, Raina E. Sacksteder, Atesh K. Worthington, Kateryna Voitiuk, Victoria T. Ly, Ryan N. Hoffman, Matthew A.T. Elliott, David F. Parks, Rebecca Ward, Sebastian Torres-Montoya, Finn Amend, Natalia Montellano Duran, Paola A. Vargas, Guadalupe Martinez, Lucia Elena Alvarado-Arnez, Drew Ehrlich, Yohei M. Rosen, Arnar Breevoort, Tallulah Schouten, Sri Kurniawan, David Haussler, Mircea Teodorescu, Mohammed A. Mostajo-Radji

## Abstract

Project-based learning (PBL) has long been recognized as an effective way to teach complex biology concepts. However, not all institutions have the resources to facilitate effective project-based coursework for students. We have developed a framework for facilitating PBL using remote-controlled internet-connected microscopes. Through this approach, one lab facility can host an experiment allowing simultaneous interaction by many students worldwide. Experiments on this platform can be run on long timescales and with materials that are typically unavailable to high school classrooms. This allows students to perform novel research projects rather than just repeat standard classroom experiments. To investigate the impact of this program, we designed and ran six user studies with students worldwide. All experiments were executed in Santa Cruz and San Francisco, California, with observations and decisions made remotely by the students using their personal computers and cellphones. In surveys gathered after the experiments’ conclusion, students reported increased excitement for science and a greater desire to pursue a career in STEM. This framework represents a novel, scalable, and effective PBL approach that has the potential to democratize biology and STEM education around the world.

## 1. Introduction

Given the rapid rise in demand for workers in science, technology, engineering, and mathematics (STEM), there is a growing need for high-quality STEM education. However, access to STEM education remains highly unequal. In the United States, Latinx people are the fastest growing demographic, currently encompassing over 19% of the country’s population while receiving less than 12% of degrees awarded in STEM fields [1, 2].

Access to quality education has further deteriorated during the COVID-19 pandemic [3]. The migration from in-person classes to remote video conferencing systems like Zoom has presented many challenges for students and educators [4]. The negative impacts of this migration have disproportionately affected students from traditionally underrepresented groups [5, 6]. This unequal learning loss, particularly in STEM disciplines, has not only widened the gap between students but has jeopardized global initiatives such as Education 2030 and the United Nations Sustainable Development Goals [7, 8]. New approaches for high-quality remote STEM education hold the potential to reverse this trend and further spread access to educational resources worldwide.

Project-based learning (PBL) is an effective approach for teaching complex STEM concepts [9, 10], particularly for students from backgrounds typically underrepresented in STEM professions [1, 11, 12, 13]. In PBL, students have the chance to learn in a hands-on matter, investigating deeper questions and discovering truth for themselves [1, 14, 13]. Despite the benefits of PBL, three significant barriers stand in the way of its broad implementation into STEM laboratory courses: 1) high local infrastructural cost required to perform complex PBL projects, 2) limited teacher training, and 3) potential exposure to hazardous materials [1, 15]. The additional challenges brought about by remote learning further compound these barriers. The lack of an effective replacement for in-person lab activities has caused sub-par learning experiences for students. Finding a way to provide students with the experience of practical project-based lab work is essential if we expect remote classes to work as well as in-person ones.

Several systems have been proposed to address this issue by facilitating at-home PBL. Different initiatives and companies have created do-it-yourself (DIY) kits that students can use to perform simple experiments [16, 17, 18, 19]. While intriguing, these kits require the acquisition and shipping of individual kits to each student, group of students, or school [20, 21] and are therefore not scalable, nor can they easily reach isolated communities. Other groups have created laboratory simulations that allow students to explore a virtual environment [22, 23], as well as tutorials and manuals for students to replicate using common household items [24, 25]. However, these solutions only enable students to explore a few canned options with a predetermined “right answer,” denying them the true experience of scientific experimentation and discovery.

It has also been shown that classroom examples that factor in local issues leads to better student engagement and learning outcomes, particularly for students from underrepresented backgrounds [1, 26, 27, 28]. This is hard for canned simulations and “kitchen chemistry” kits to do. Therefore, an approach that is scalable at a low cost, adaptable to local contexts, and easily accessible to students has the potential to revolutionize STEM education by democratizing PBL implementation in classrooms globally.

The Internet of Things (IoT) has transformed many fields of society and research, including agriculture [29], healthcare [30] and wildlife conservation [31]. Yet, the adoption of IoT in the classroom has been minimal [32]. When applied to the remote operation of lab equipment, IoT can be used to create educational tools. Early work using internet-connected single camera microscopes used photophobic organism samples and enabled remote control of light stimulation to create interactive experiments [33, 34]. The activities enabled by these devices are short and generally designed to fit in the time frame of a single class session. Programs like this provide a proof of principle for the usability of IoT technologies in education. However, these technologies are not designed to be adaptable for context-informed PBL and running experiments requiring multiple conditions over longer multi-day periods. One recent study used an IoT system in a longer-term experiment. Students remotely monitored soil moisture while evaluating the effects of ground cover on plant seedlings [35]. However, this project was not fully remote and required students to make measurements in person at the site of the experiment, and must therefore be confined to a defined geographical location.

When considering the technologies needed to set up a fully remote, longer-term experiment, remote microscopy is alluring in its simplicity. Simply point a camera at a phenomenon, and people can make observations from anywhere. Existing microscope systems can be modified with widely available internet-connected cameras, or purpose-built connected imaging systems can be developed. Unlike systems that involve direct physical interaction with samples, remote imaging systems are non-invasive and capable of being set up at a low cost. Imaging systems capable of performing near real-time simultaneous longitudinal tracking of cells, 3D cultures, and small organisms have started to emerge in research settings [36, 37, 38]. These systems can be used to perform experiments with a diversity of models and conditions. The rapid cloud-based transfer and storage of data and their relatively simple interfaces make them excellent tools for science [37, 38]. Given the versatility of these devices, applying them to remote education represents a cost-effective and scalable approach to performing PBL.

Here we describe the implementation of these technologies in the biology classrooms of Latinx communities in the United States and Latin America using examples relevant to their local surroundings. We demonstrate the capability of this framework to perform a single shared experiment for several groups around the world simultaneously. Furthermore, we provide a framework for interested parties to remotely perform their context-informed PBL-based courses.

To evaluate this program’s utility, scalability, and impact, we ran several iterations of the PBL-based remote class with different user groups. We conducted user surveys with two of the participant groups: high school students in California’s Central Valley and college students in Bolivia. We compared the user experiences of both groups and contrast them with previous work performing in-person PBL with similar cohorts [1]. We show that IoT-enabled remote PBL is an effective and scalable approach for serving underrepresented students in STEM and provides a novel platform for comparative experimental STEM education studies.

## 2. Program Development

### 2.1. Designing a Curriculum

We defined a roadmap for designing a curriculum program using a remote microscopy system to facilitate project-based experimental biology. The roadmap is aimed at research groups interested in using their lab to facilitate a remote PBL program. The program is designed as a supplement to existing biology courses, facilitators would be researchers from the lab that wish to participate in educational collaboration with a teacher’s class. In the weeks leading up to the remote experiment, and facilitators teach supplemental lessons on various scientific topics. The lessons build on the student’s current knowledge and prepare them to design their biological experiment. After the experiment, students analyze their findings and present a conclusion to the class. A broad outline of this framework is shown in Figure 1.

**Figure 1:**
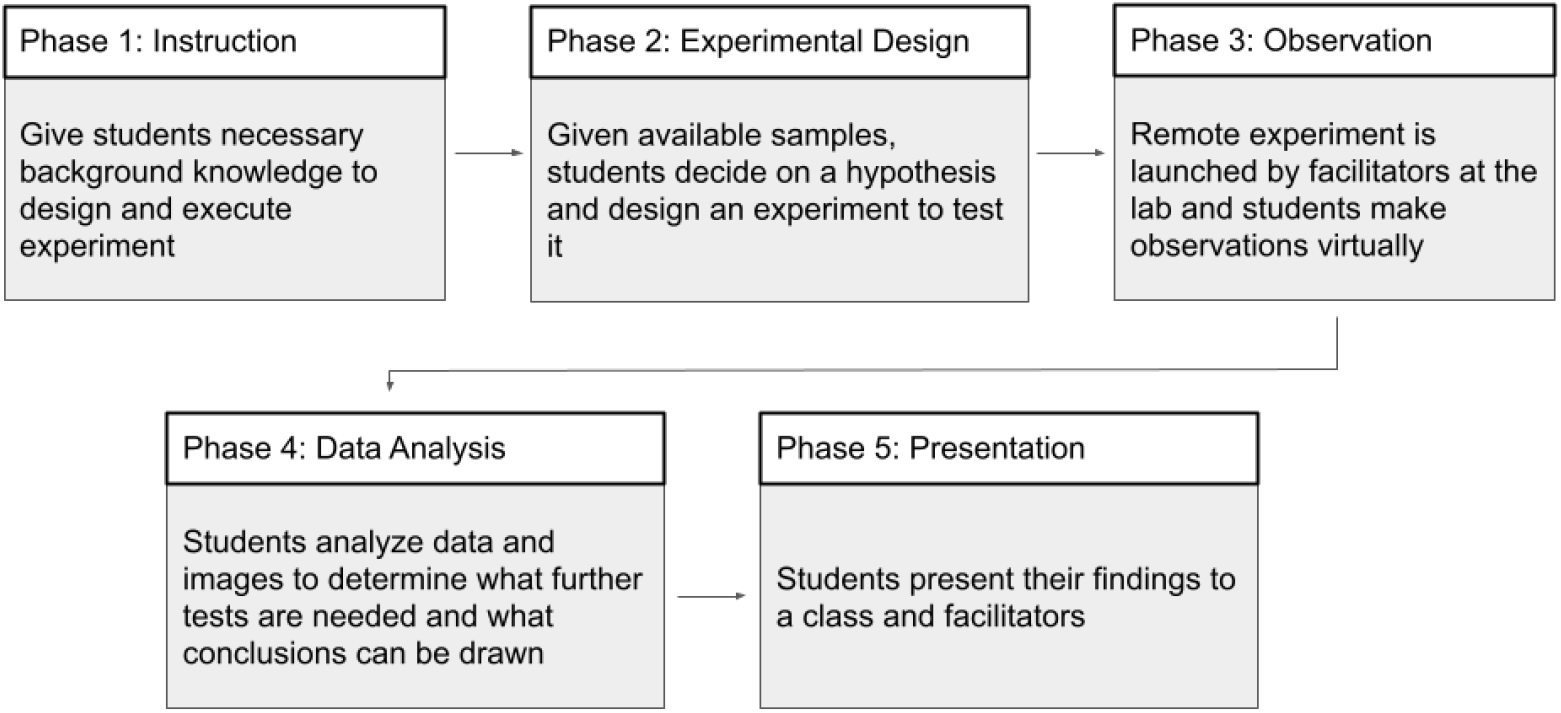
Program Roadmap.

#### 2.1.1. Phase 1: Instruction Modules

The first phase of this remote experimental biology course is to provide students with the instruction to support them through designing their experiments, collecting and analyzing data, and coming to scientifically supported conclusions. This is done through a series of lectures and activities in the weeks leading up to the experiment.

Developing a lesson involves first identifying the desired learning outcomes. Desired learning outcomes describe what students should know or be able to do at the end of a lesson. It is useful to clearly articulate the learning outcomes to help inform the students of what to expect from the lesson and gauge their learning. Additionally, they guide the instructor in lesson design. Designing a lesson should begin with articulating 2-3 learning outcomes. Resources on how to design learning outcomes can be found at [39, 40].

Rather than replicating the content the students are already learning in class, facilitators should aim to show students relevant topics from the perspective of a researcher, with a focus on how to approach a subject from an experimental design standpoint. Example activities from lessons we ran can be found in the Supplemental Materials section.

#### 2.1.2. Phase 2: Experiment Design

After facilitators have run lessons presenting the necessary background knowledge, it is time to design the experiment. A good experiment for our platform is one where the sample requires no maintenance after setup, and the observations of interest are all visual. Facilitators introduce the available biological samples and then guide students through defining a testable question and designing an experiment to address that question. The students will then create a proposal outlining their question, experimental design, and hypothesis to be tested. An example activity to help students propose their hypotheses and plan their experiment can be found in the Supplemental Materials section. With this proposal, the project facilitators will prepare the appropriate samples and set up the remote imaging device. After the setup, the experiment is launched.

#### 2.1.3. Phase 3: Observation

Next, students make observations as the experiment runs in real-time. The most substantial hurdle to conducting a successful remote biology course is providing students with the experience of real-time observations of an experiment that does not have a predetermined outcome. This is the essence of doing science, so this issue must be faced. Once the experiment is launched, the device is set up in such a way as to allow students to make observations from their own cell phones or computers at any time, and they may do so on nights or weekends when there is no facilitator to help them interpret what they are seeing happening in the biological system These real-time observations are analogous to looking directly through a microscope. Students should capture and log their observations for later use, i.e. keep an elementary lab notebook.

#### 2.1.4. Phase 4: Data Analysis

Finally, the students compile their observations and analyze their data. In addition to live monitoring, time-stamped historical data is also accessible. This allows students to revisit earlier time points to make direct measurements for comparative analysis or make additional observations. Data analysis and the related computer skills required for this are a critical component of modern scientific research that is typically under-emphasized in early college biology coursework [41]. By providing students with simple-to-use analysis tools, students are empowered to look closer at what is occurring and draw more interesting conclusions. For an image-based experiment, this can be accomplished with software that allows students to annotate times-tamped images in the data set such that features can be visually tracked over time, and students can note observations on them (Figure 2).

**Figure 2:**
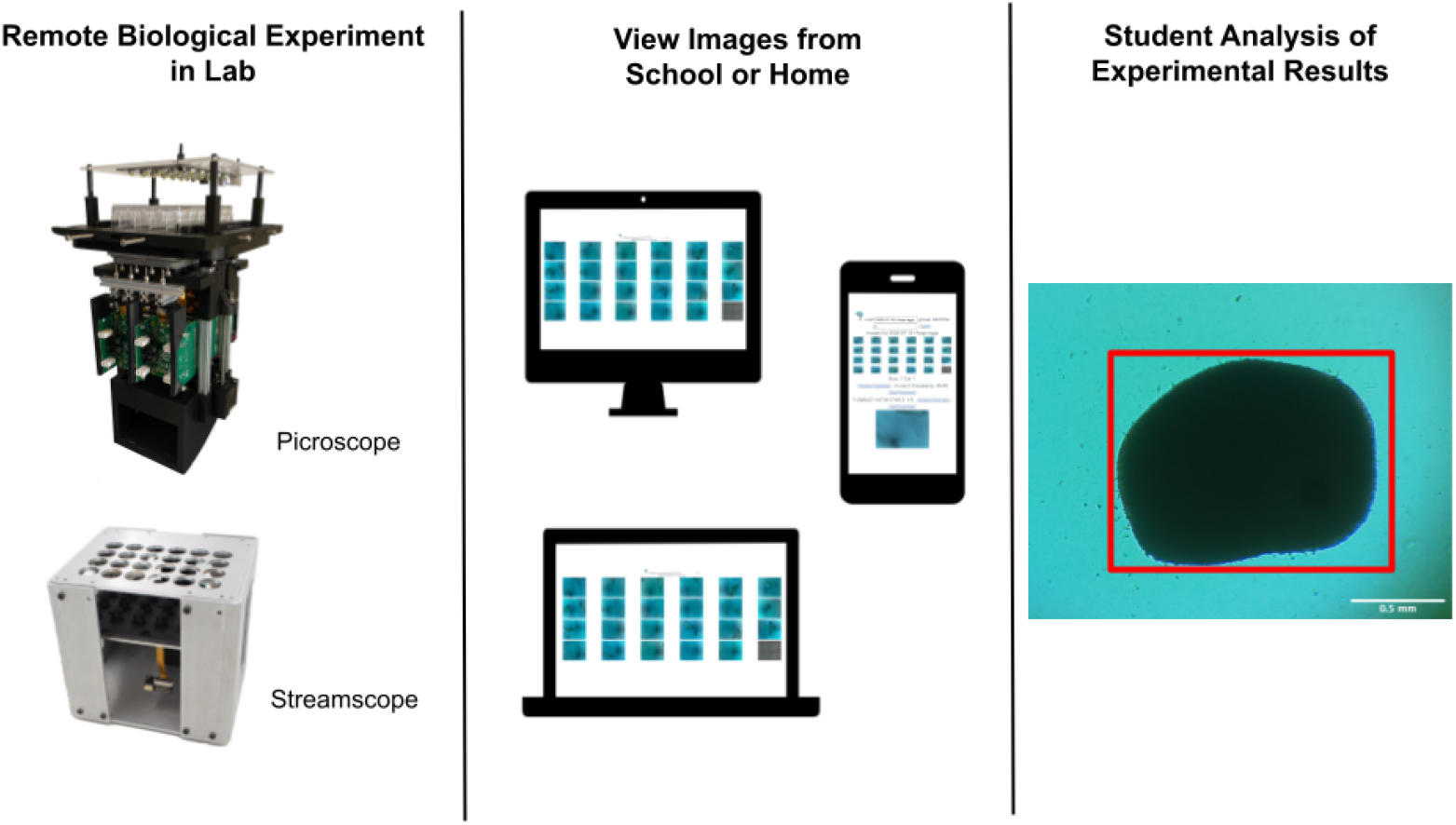
Remote experiment workflow. Data is recorded in the remote lab using the Pircoscope or Streamscope, viewed from personal devices, and analyzed by the students.

#### 2.1.5. Phase 5: Presentation

When the experiment has concluded and the data has been analyzed, the students present their findings. They should offer a scientific conclusion based on the data and then defend their analysis against questions asked by the instructors and their peers. Presentations can come in many forms, including live slide show presentations, video production, or written reports. An example scientific presentation outline is provided in the Supplmental Materials section. This mirrors how new science is reported by professionals working in the field and challenges the students to consider the most effective ways to communicate science.

## 3. Methods

### 3.1. Imaging System Selection

Any IoT-based imaging for remote experimental education needs to allow the capture of longitudinal data from the culture plate on an appropriate timescale for the samples. The system should make this data readily available for the students and other users, ideally allowing them to access it from their own devices. Finally, the data should be easily interpretable and permissible for analysis. In the six studies we ran (further described in section 4), we used a system called the “Picroscope” in all of them and add another system called the “Streamscope” in two of them.

The “Picroscope” is a low-cost open-source IoT-enabled remote microscopy device built primarily using off-the-shelf components and 3D printed pieces [36, 37] (Figure 3). The Picroscope captures 3D z-stack data on long timescales and makes the image datasets available to students on an image viewer website (described in Supplemental Video 1). It is designed to image cells, tissues, and organisms growing in the wells of a plastic plate. Picroscopes can also be deployed in a standard temperature and humidity-controlled CO_2_ incubator, allowing the use of biological cultures and samples that require that kind of controlled environment to thrive. The “Streamscope” is another maker microscope that live-streams video from the wells simultaneously. Students view the live stream through YouTube. The 3D z-stack image data from a Picroscope is well-suited to samples that are adhered to a well plate where growth is effectively visualized on the scale of hours and days. For faster moving samples like zebrafish or other small animals, the z-stack functionality is less useful, and users benefit more from viewing a video stream. It is important to select the right system for the samples available to you or to select appropriate samples for the imaging system you have.

**Figure 3:**
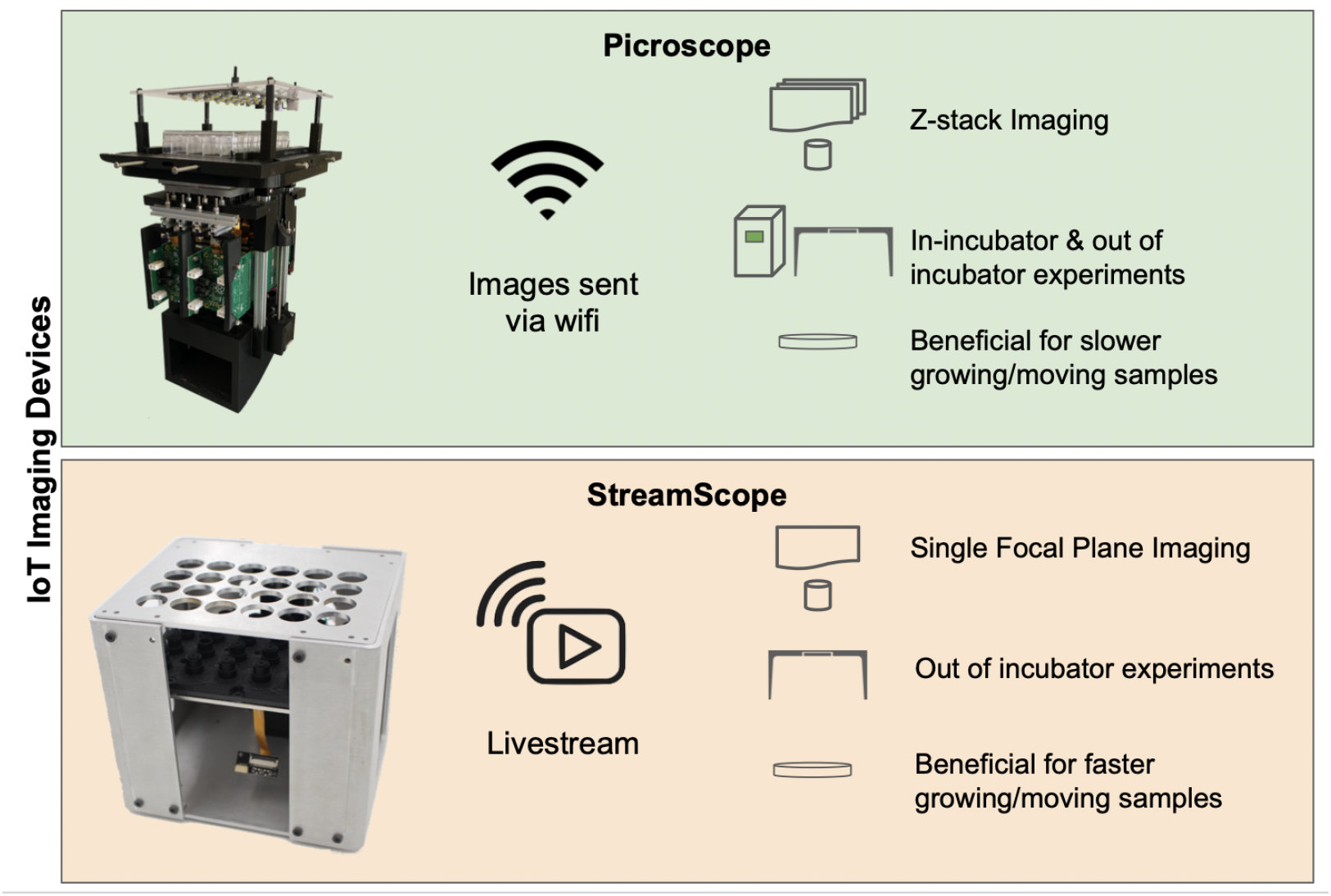
Comparison of remote imaging devices used in our program.

The Picroscope contains 24 cameras imaging in parallel, allowing 24 students to each capture image data from their own well or all students to view all 24 wells. The wells are formatted into a standard 24-well cell culture plate. Images are uploaded to the internet, where they can be viewed through a public website. Uploads occur immediately after a timestep is captured, giving end users a near real-time experience looking through a virtual microscope. These image datasets are also stored long-term, allowing for experiments to be replayed or loaded into data analysis pipelines. Picroscope experiments can be run inside a standard CO2 and humidity-controlled incubator or outside, giving experiment designers flexibility with the samples they choose.

#### 3.2. Data Analysis Pipeline

Since high school and early college-level biology classes don’t typically have a programming requirement, it is important to provide user-friendly tools for analysis. A balance should be struck between introducing students to working with code and not requiring them to write much code themselves.

In the programs we ran, students completed a learning module on an online Jupyter notebook server [42] where they learned the basic fundamentals of manipulating experimental data in Python. The module provides them with a coding environment, example code, and online video tutorials that they can follow. A facilitator lectures the students and guides them through completing the learning module. At the end of the experiment, students are guided through the analysis of the data they have collected throughout the experiment and then give a presentation on their findings. Microscopy images recorded from the experiment are analyzed on the notebook server, where a Python GUI application for annotating images was created so that students can highlight findings that they consider significant throughout the experiment. For example, students can classify cancer cells and annotate specific phenotypes they believe are changed by the application of drugs.

#### 3.3. User Studies

Student users were selected through partnerships with different organizations. Students in California were 4th year high school students at Alisal High School, located in Salinas, California. All students were part of an Advanced Placement (AP) Biology course.

Students in Bolivia came from two different universities:

1. First-year students in the Biotechnology degree at the Universidad Catolica Boliviana San Pablo in Santa Cruz de la Sierra, Bolivia, were the students that performed the experiments in neuroblastoma cell lines.
2. Second-year Biochemistry students at Franz Tamayo University performed the studies of chlorine dioxide. Franz Tamayo University has 4 campuses in Bolivia: Santa Cruz de la Sierra, La Paz, El Alto and Cochabamba. The students were from all campuses.

Students for the multinational study involving Bolivia, Mexico, Peru, Brazil and Spain were students undertaking a two weekend-long course through the nonprofit organization Science Clubs International.

#### 3.4. Remote Teaching

Before each course started, we met with the lead instructor and designed the experiments that were relevant to the course. When applicable, we retained the core curricula from the national standards. All supplemental teaching by UCSC researchers was facilitated using Zoom. At Alisal High School, classes took place once a week. For the Science Clubs International, the courses were taught in two sessions, one week apart from each other. For the university courses, the classes were taught over a three-week period. Instructors were graduate students at the University of California, Santa Cruz. The courses in Bolivia were taught in Spanish, while the courses in the United States and through Science Clubs International were taught in English.

#### 3.5. Surveys and Subject Tests

Surveys were conducted using Google Forms; all questions are listed in Supplemental Table 1.

Students at the Catholic University of Bolivia were tested in person using a subject test designed by the course lead instructor. The translation of the test can be accessed in Supplemental Note 6.

#### 3.6. Statistical Analyses

Statistical comparisons between groups of students were conducted using the Mann-Whitney test. Tests of scores were analyzed using the Student’s t-test. Significance was established as follows: * =*p* < 0.05; ** = *p* < 0.01; *** = *p* < 0.001; **** = *p* < 0.0001.

#### 3.7. Culture of Neuroblastoma Cells

Mouse neuroblastoma cells (ATCC # CCL-131) were cultured in standard tissue culture incubators at 37°C in Dulbecco’s Modified Eagle’s Medium -high glucose (Millipore Sigma # D6429) supplemented with 10% Fetal Bovine Serum (Thermo Fisher Scientific # 26140079).

Drugs used were Retinoic Acid (10 *μ*M, Millipore Sigma # R2625), Neurodazine (2 *μ*M, Millipore Sigma # N6664), Primocin (2 *μ*l/ml, Invivogen # Ant-pm-05).

#### 3.8. Zebrafish experiments

Fertilized *Danio rerio* eggs were purchased from Carolina Biological Supply Company (Catalog # 155591) and maintained in media containing 15 mM sodium chloride (Millipore Sigma # S9888), 0.5 mm potassium chloride (Millipore Sigma # P3911), 1 mM calcium chloride dihydrate (Millipore Sigma # 223506), 1 mM magnesium sulfate heptahydrate (Millipore Sigma #1058822500), 150 *μ*M potassium phosphate monobasic (Millipore Sigma #P5655), 50 *μ*M sodium phosphate dibasic heptahydrate (Millipore Sigma #S9390), 0.7 mM sodium bicarbonate (Millipore Sigma # S5761), and 0.1% methylene blue (Millipore Sigma #M9140).

The following drugs were used in the different user studies: caffeine (10 *μ*M, Millipore Sigma # C0750), ammonium nitrate (10 mg/L and 100 mg/L, Millipore Sigma # 221244), (-)-Nicotine (1%, Millipore Sigma # N3876), and chlorine dioxide (0.1%, 0.5% and 1%, Novatech # R-8039A). Gold and graphene nanoparticles were synthesized in-house and used at concentrations ranging from 10 to 100 nM.

## 4. Results

### 4.1. Remote Experimental Instruction Allows for Scalable and Adaptable PBL in Underserved Communities

We ran four experiments with different user groups to evaluate the flexibility of our framework and possible experiments we can run.

As context-informed education has been shown to be particularly effective [1], we coordinated with local teachers to ensure that the projects related to real-world problems of interest to the students. These experiments were performed in San Francisco, CA, while students accessed the data in near real-time from their homes. Experimental details can be found in Table 1.

**Table 1:** Summary of user studies performed.

| Experiment Theme | Biological Sample | Experiment Condition | Number of Users | Geographical Area | Distance from the experiment |
| --- | --- | --- | --- | --- | --- |
| Effects of caffeine on development | Zebrafish embryos | Caffeine | 12 | Salinas, CA, USA | 170 Km. |
| Effects of agriculture byproducts on development and physiology | Zebrafish embryos | Ammonium nitrate, nicotine and caffeine | 20 | Salinas, CA, USA | 170 Km. |
| Biocompatibility of nanoparticles | Zebrafish embryos | Gold and Graphene Nanoparticles | 20 | Bolivia, Mexico, Peru, Brazil, and Spain | Up to 9,700 Km. |
| Toxicity of chlorine dioxide | Zebrafish embryos | Chlorine dioxide | 131 | Bolivia: Santa Cruz de la Sierra, Cochabamba, La Paz, and El Alto | Up to 8,700 Km. |

#### Study 1: Alisal High School

As caffeine consumption is higher in student groups compared to the general population [43], we focused our first study on the effects of caffeine exposure on organism development. Students in the AP Biology course at Alisal High School in Salinas, CA observed the effects of three different caffeine concentrations in developing zebrafish embryos. Observations included effects in the whole body, such as movement and twitching, and effects on specific organs, such as the heart.

#### Study 2: Alisal High School

A second user study with a new cohort of AP Biology students in Alisal High School focused on the effects of common byproducts of agricultural activities. We focused on three chemicals: 1) Ammonium nitrate, a chemical commonly used in fertilizers [44]; 2) nicotine, historically used as a pesticide and being the chemical basis of several modern neonicotinoid pesticides [45, 46]; and 3) caffeine, a contaminant in cultures of several plants and crops [47]. The students observed the effects of these chemicals in the developing zebrafish embryo. Descriptions of their observations unveiled novel discoveries, such as a delay in fin development in zebrafish exposed to ammonium nitrate. Students made new discoveries, carefully measuring for the first time the effects of different levels of ammonium nitrate on zebrafish embryonic development.

#### Study 3: Five Countries

While there are variable degrees of Internet reliability throughout the United States, its adoption is relatively high compared to the rest of the world [48]. To understand if places with less reliable internet could adopt IoT-enabled PBL, we performed a user study with 20 students in five countries: Brazil, Bolivia, Spain, Mexico, and Peru. The study complemented an online outreach activity by a United States-based non-profit organization Science Clubs International, which targets high school and early college students in the named countries. The students used remote microscopy to perform biocompatibility studies of custom-made gold and graphene nanoparticles and determine the optimal concentration of such particles to be used in the bioengineering context.

#### Study 4: Bolivia

Finally, we focused on the scalability of our approach. We performed a 4th user study complementing a pharmacology college-level course in Bolivia. This course had over 130 students enrolled across four cities in the country: Santa Cruz de la Sierra, La Paz, El Alto, and Cochabamba. Bolivia has the slowest Internet connection in South America, which makes remote education challenging. Indeed, the government and the education sector have often turned to television and radio as means to conduct large-scale remote education in situations such as the COVID-19 pandemic [49]. In this study, the students tested the toxic effects of chlorine dioxide in animal physiology.

Chlorine dioxide is a type of industrial bleach that became highly popular in Latin America during the COVID-19 pandemic, as several politicians promoted its use to prevent SARS-CoV-2 infection [50].

Altogether, these user studies allowed us to conclude that remote experimental biology PBL can be effectively adapted for the needs of different groups; that it can be used to reach underserved communities; and that it is scalable to reach a large number of students in the same or different countries simultaneously.

### 4.2. IoT-enabled PBL Positively Affects Different Latinx Populations

To understand how our approach impacted different populations of Latinx students, we performed user studies with two distinct student groups: 1) High school students in the AP Biology course at Alisal High School in Salinas, California, and 2) First year General Biology students at the Catholic University of Bolivia in Santa Cruz de la Sierra, Bolivia. Both Salinas and Santa Cruz de la Sierra are agricultural hubs in their respective regions, and both have a population with a significant Latinx majority [51, 52].

Due to their curricular equivalency, in the United States, AP Biology courses are used to substitute first-year college-level General Biology courses [53]. On the other hand, the Bolivian education system does not have an AP Biology equivalent course in high school, and students are first exposed to complex concepts in cellular and molecular biology in their first year of college. Therefore, we designed and implemented a similar PBL approach for both groups. Given that agriculture is the major economic driver in both regions, the project was designed to address a common issue in fertilizer exposure: the development of childhood cancer [54]. Specifically, neuroblastoma has been strongly associated with children raised in agricultural areas [55]. Leading up to the experiment, we lead additional lectures for both cohorts on various topics, including cell signaling, development, microscopy, data analysis, and the scientific method (Table 2). Complete lesson outlines, learning outcomes, and some examples of activity materials for each lesson can be found in Table 2 and Supplemental Notes 1-5. The students were then given a project in which they had to investigate the effects of three drugs in neuroblastoma cell lines using an IoT-enabled microscope. The three drugs selected were: Retinoic acid, a vitamin A-derivative that is an effective treatment for high-risk neuroblastoma [56]; Neurodazine, a synthetic drug that is known to differentiate neuroblastoma cells into neurons [57]; and Primocin, a proprietary mixture of four antibiotics and antimycotics that is commonly used in cell culture to prevent bacterial and fungal contamination due to its low toxicity to mammalian cells [58]. The experiment was set up using a Picroscope in a lab in Santa Cruz, CA. The students analyzed the effects of each drug and a no-drug control every hour for a period of 5 days. They split into groups and tracked individual cells’ migration, division, differentiation, and survival. Groups then discussed their results with their peers and reached similar conclusions: the students found a strong effect in cell proliferation and survival upon retinoic acid treatment, an effect in morphology upon neurodiazine treatment, and no effect of primocin in neuroblastoma.

**Table 2:**
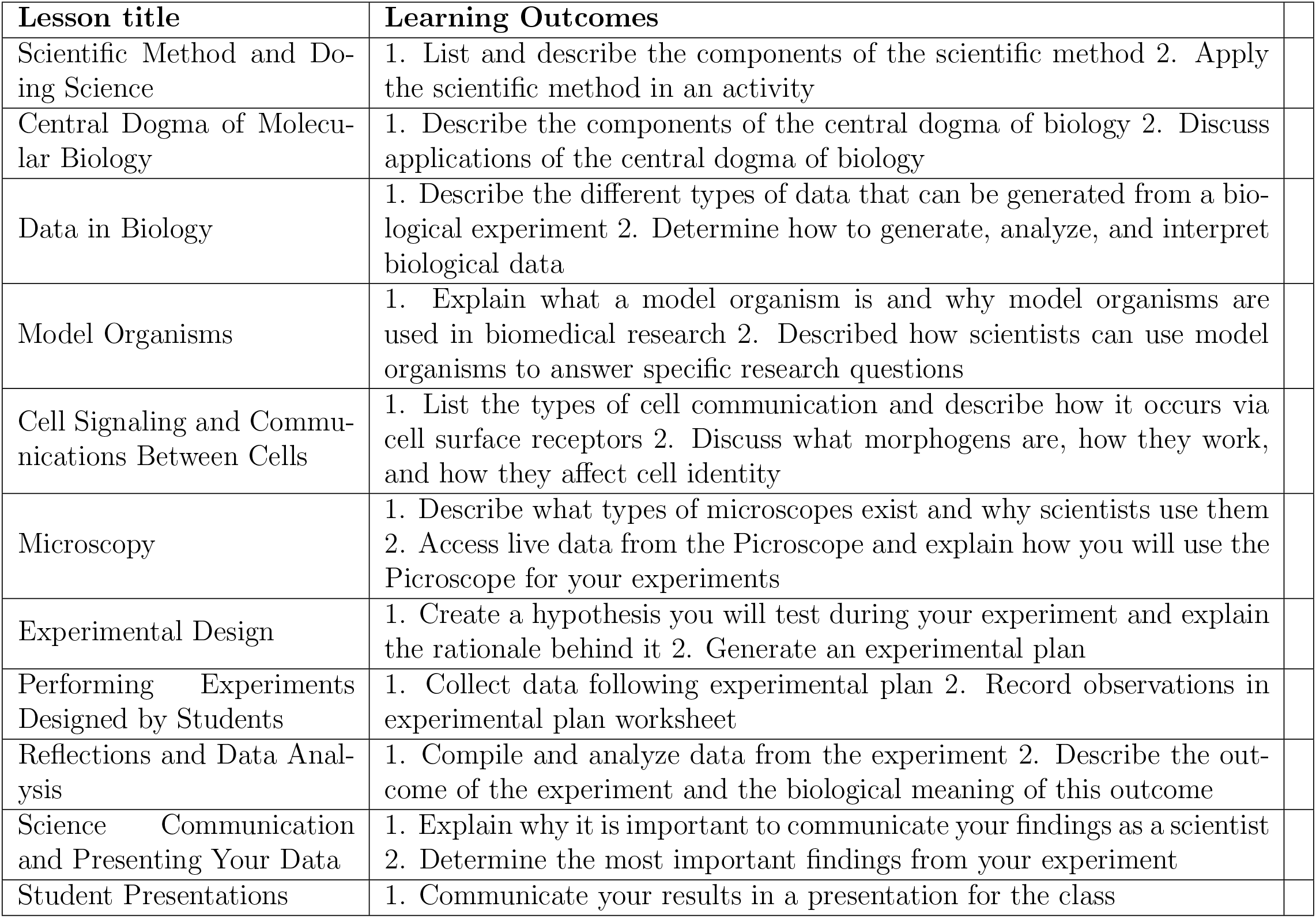
Lesson Plans.

The students then presented their data in different formats: The California-based students recorded videos to disseminate their discoveries over social media (Supplemental Video 2), while the Bolivia-based students presented their results at their university science fair (Supplemental Figure 1).

After completing their respective courses, we anonymously surveyed the students with two different instruments for quantifying STEM identity: The STEM Professional Identity Overlap (STEM-PIO-1) and the Role Identity Survey in STEM (RIS-STEM). STEM-PIO-1 is a single-item survey that measures the self-perceived overlap of students with STEM professionals [59]. RIS-STEM is a 26-question survey designed to measure several aspects of STEM identity, including competence in STEM, interest in STEM, self-recognition in STEM, and recognition by others in STEM [60]. We then asked the students how participating in our remote PBL program affected their answers to the STEM identity questions.

Responses on the STEM-PIO-1 (Figure 6) indicate similar self-perceived overlap with a STEM professional between the two groups (p >0.05). These results are comparable to those of students in STEM majors, as reported previously [59]. When surveyed, the majority of students in both groups reported that participating in this program had a positive effect on their STEM-PIO-1 responses.

**Figure 4:**
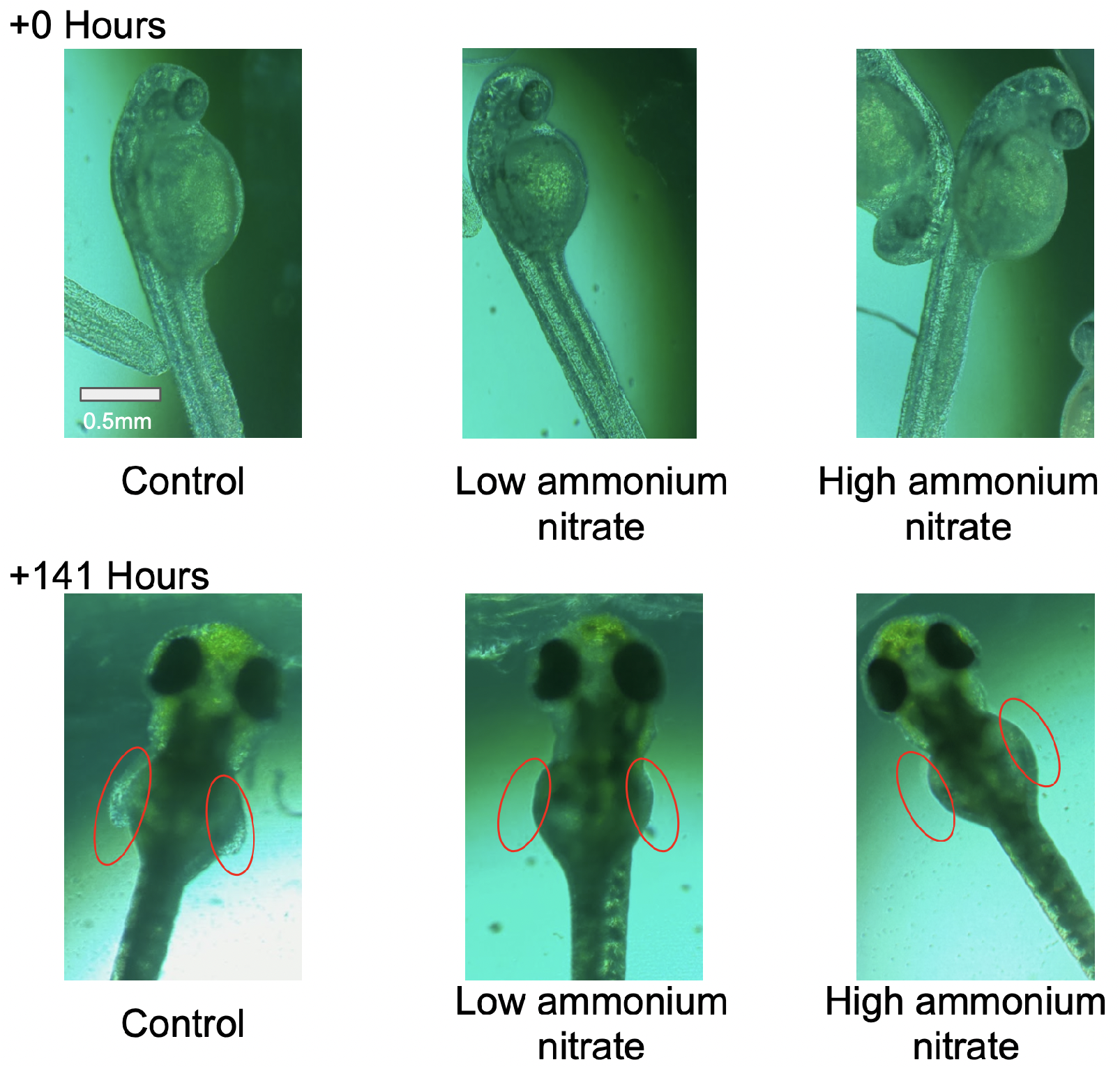
Context-informed PBL using whole organisms. Students tracked the effects of chemicals, such as ammonium nitrate, in the development of zebrafish embryos. Representative images over a 141 hours show that ammonium nitrate affects fin development.

**Figure 5:**
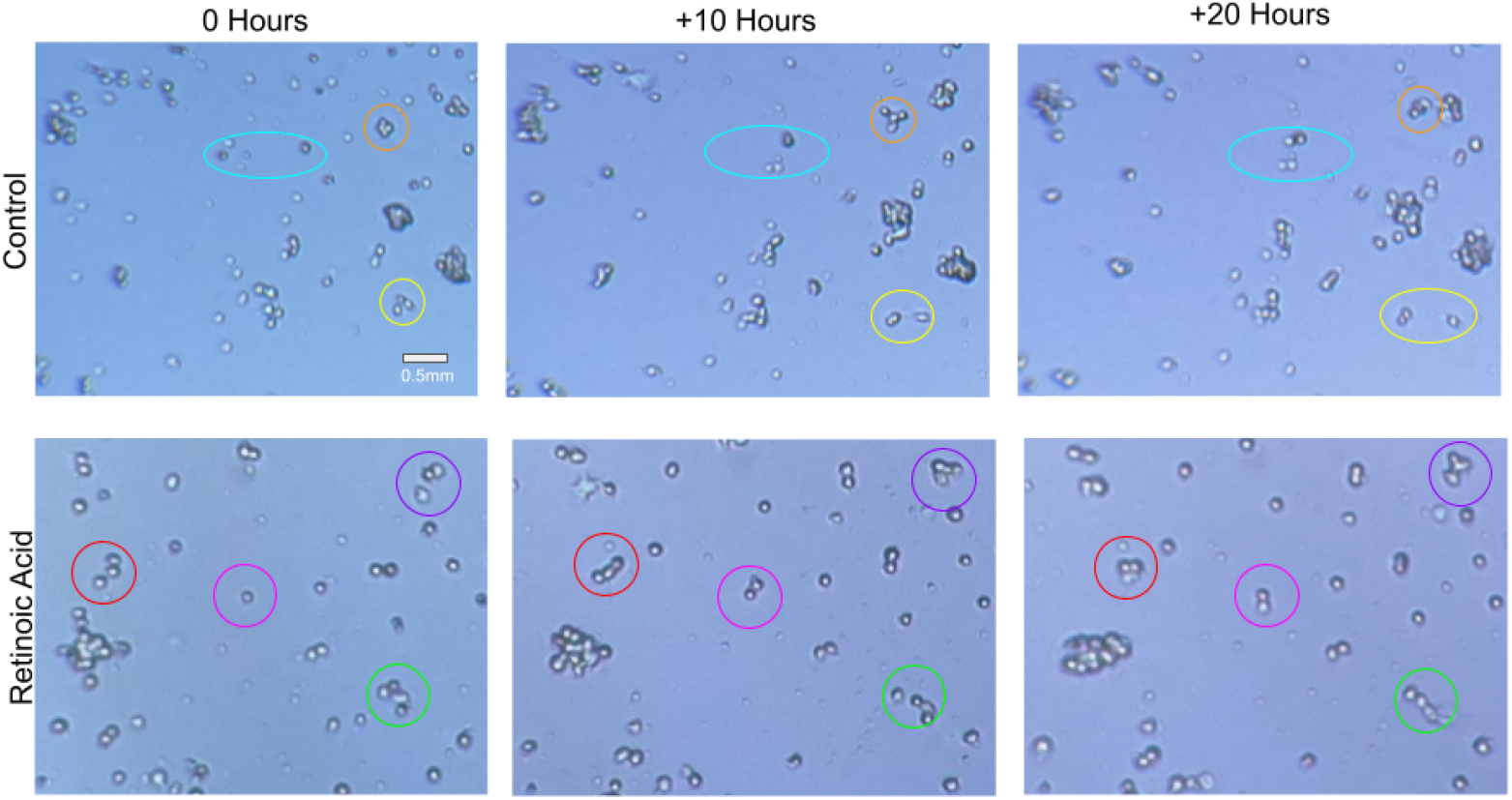
Integration of mammalian cell culture experiments in the classroom. The students used IoT-enabled microscopy to understand the effects of drugs, such as retinoic acid, in neuroblastoma cells. Representative images show the tracking of individual cells and cell clusters over 20 hours.

**Figure 6:**
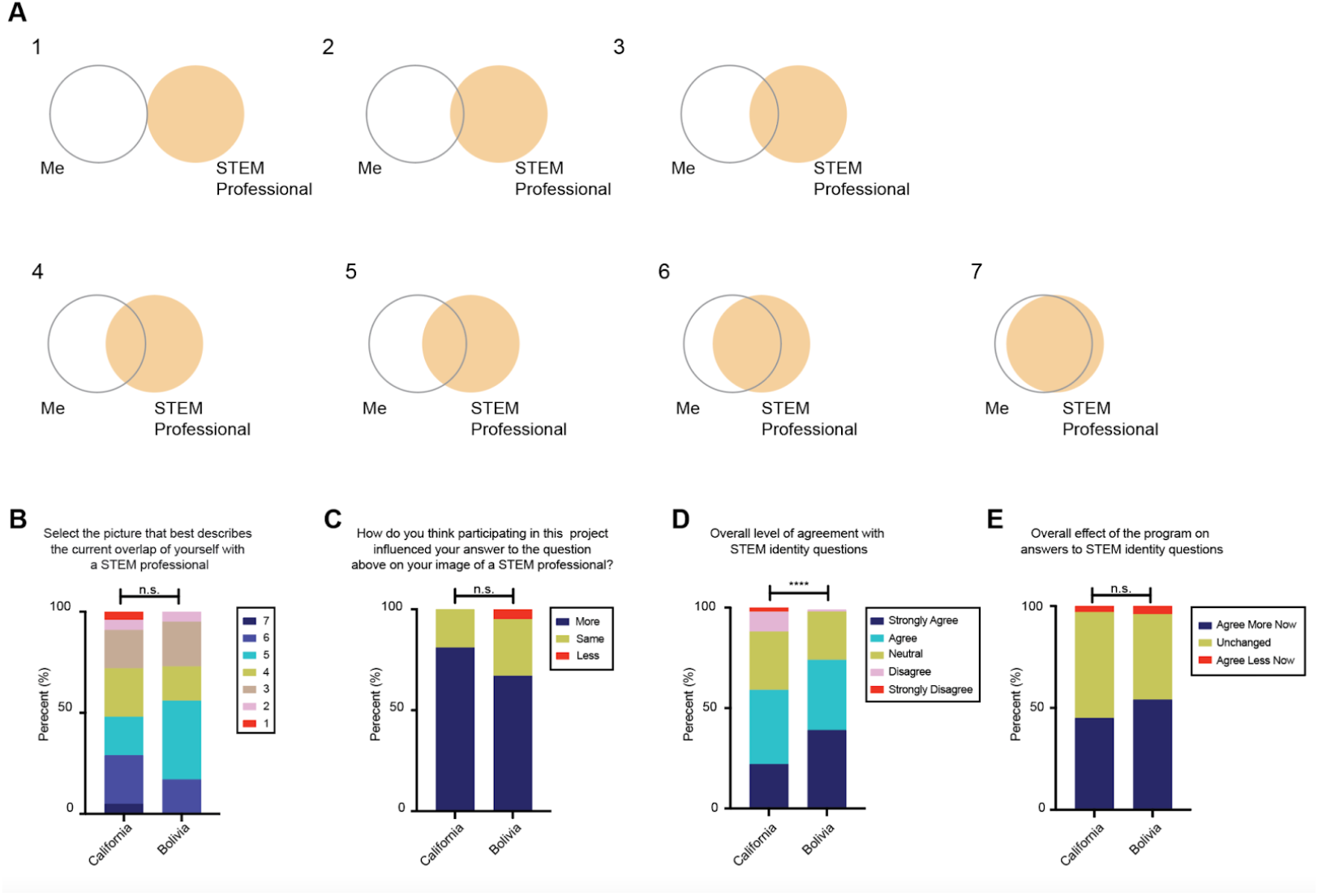
IoT-enabled PBL positively affects STEM identity in different Latinx cohorts. (A) Diagrams in the STEM-PIO-1 instrument used to assess STEM identity. Students are asked to select the picture that best describes their overlap with a STEM professional. (B) Level of agreement with the question “Select the picture that best describes the current overlap of yourself with a STEM professional”, referring to the image in A. (C) Impact of the remote instruction project on the answer to the question in B. (D) Overall level of agreement with 26 questions to assess STEM identity in the RIS-STEM instrument. (E) Overall impact of the remote instruction project on the level of agreement with statements in D. Cohort sizes: California n = 21, Bolivia n = 18. Mann Whitney test. **** = p <0.0001; n.s. = not significant.

The Bolivia students agreed significantly more with RIS-STEM questions (Figure 7) that involved doing and learning more about STEM in the future compared to the California students. For instance, 56% of Bolivia students selected “strongly agree”to “I want to learn as much as possible about STEM”, compared to only 14% of California students (Mann Whitney test, p = 0.0038) (Figure 7). The remaining 20 RIS-STEM statements received similar answers from both cohorts (Supplemental Figure 2).

**Figure 7:**
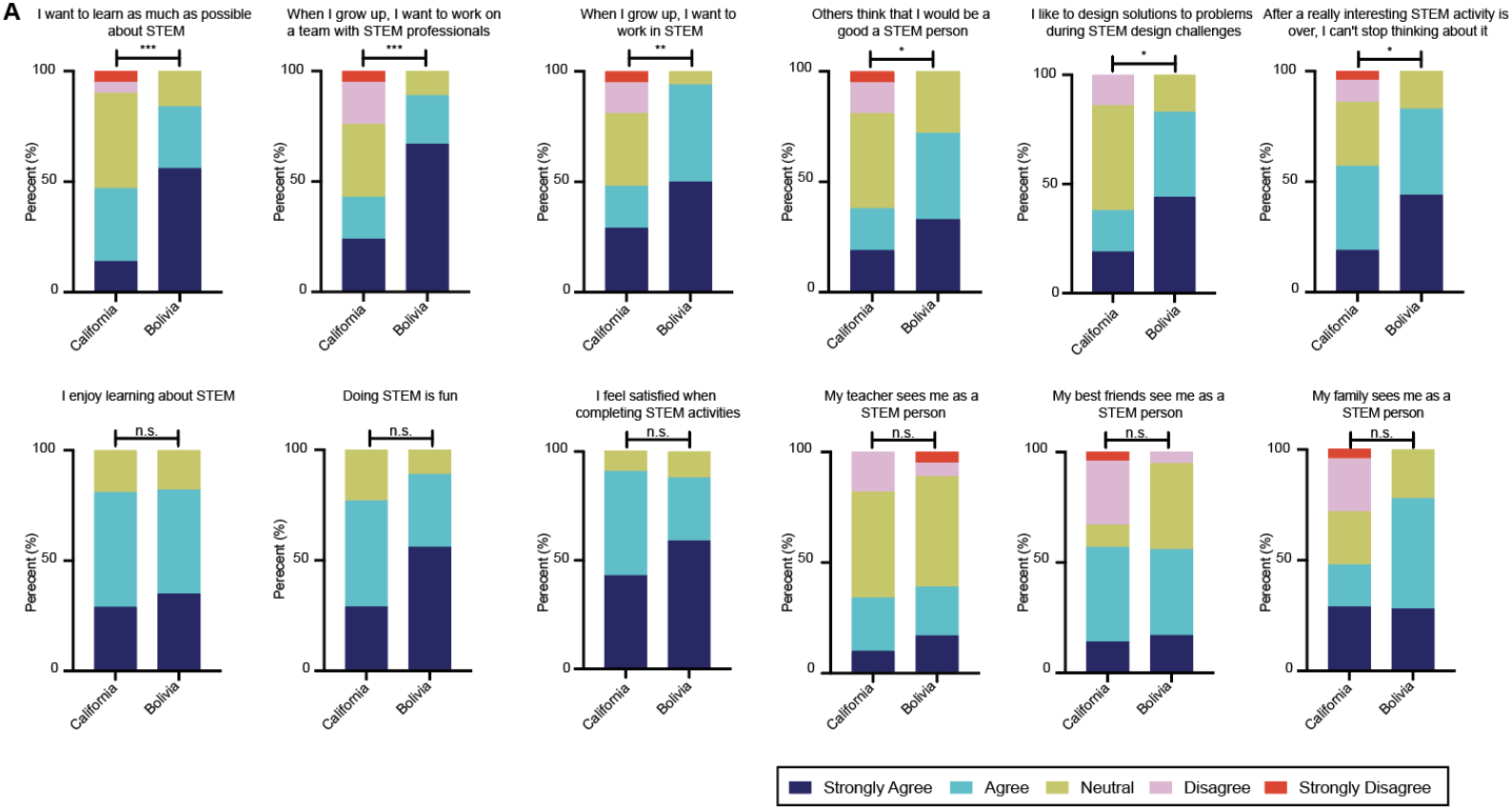
Different Latinx student groups have distinct STEM identities. (A) Students’ answers to the questions in the RIS-STEM instrument. Top: Statistically significant answers between student cohorts in California and Bolivia. Bottom: Sample statistically not significant answers between Californian and Bolivian students. Additional answers in Supplemental Figure 1. Cohort sizes: California n = 21, Bolivia n = 18. Mann Whitney test. *** = p <0.001; ** = p <0.01; * = p <0.05; n.s. = not significant.

Despite the differences in their agreements with the STEM identity questions, both California and Bolivia students had similar answers to how their participation in the project affected each of their selections (Supplemental Figure 3). Together, these results indicate that both groups enjoy learning about and doing STEM and have similar perceived support but exhibit differences in desire to pursue a career in STEM. Still, our remote PBL program positively impacted STEM identity within both cohorts, regardless of differences in STEM identity or career pursuits.

### 4.3. Student Comments Indicate Positive Experiences With Our Remote PBL Program in Both Latinx Cohorts

Beyond the multiple-choice sections of the survey, students were also given questions with free response answers. The ability for the students to easily use the imaging viewer is crucial for them to observe and analyze the results of their experiment. In the survey, we asked the students for feedback on the imaging viewer, and received overwhelmingly positive results:

> *“The imaging viewer was very easy to access and after being taught on how to understand it, it was very easy to look for the data we wanted and find clear pictures to present.”*
>
> *(California Student)*

> *“The imaging viewer was very user-friendly. I had no trouble clicking back and forth through pictures.”*
>
> *(California Student)*

> *“It was very easy to use the website to access the images.”*
>
> *(Bolivia student, translated from Spanish)*

We also asked the students whether they enjoyed being able to watch a live experiment in near real time, and if so, what they liked about it. The students largely appreciated the live aspect of the experiment:

> *“Yes, I did enjoy watching a live experiment in almost real time because it felt like it was in a real lab and I was able to see how it was that an experiment is set up.” (California Student)*
>
> *“It was really interesting being able to come back a day later to see how much the cells had changed and making discoveries in real time rather than looking at something that’s already been observed in the past.” (California Student)*
>
> *“I loved it! It was very interesting to run a long-term project, as we had the opportunity to constantly collect data and progressively get new results that we could see in real time.”*
>
> *(Bolivia student, translated from Spanish)*
>
> *“I liked being able to work with samples that are thousands of kilometers away, I feel that it is something that opens the doors for us to have a more entertaining and broader education.”*
>
> *(Bolivia student, translated from Spanish)*

One of the goals of the project is to increase students’ knowledge and skills in scientific research. To achieve this goal, students were exposed to techniques in experimental design, data collection, and data analysis throughout the course and in their final project. When asked to report “the most interesting or useful thing you feel you learned in a post-course evaluation”, several scientific skills were mentioned:

> *“The most useful thing I feel I learned was knowing how to use a microscope and collecting data properly.”*
>
> *(California Student)*
>
> *“The most useful thing I learned was how to analyze the data and how to practice coding.” (California Student)*
>
> *“I learned how to collect data and apply it to a real situation.”*
>
> *(California Student)*

The students also enjoyed the visual nature of the project, with one student writing:

> *“The most interesting thing I learned was how to visually see and recognize normal cells and cancer cells. Most useful thing I learned was how to properly analyze an image.”*

Their interest in the project topic, cancer cell growth, was also commonly mentioned throughout the students’ evaluation responses. Some more examples of their responses to the same question as above, “What was the most interesting or useful thing you feel you learned?” included:

> *“The most interesting thing I learned was how cancer cells look over time because I didn’t know how they looked before.”*
>
> *(California Student)*
>
> *“The most interesting thing I learned was about the three different types of medicines that can be used against cancer cells and how those affected the growth of cancer cells, it was also interesting to look over the images of the cancer cells and see them close up to analyze them.” (California Student)*
>
> *“I feel like I have better knowledge about cell functions and how different factors can affect the body. Going into depth about cell death and growth was mind-blowing and made me much more aware of what happens in our body.” (California Student)*
>
> *“How neuroblastoma cells behave with certain drugs and how that can be used to improve people’s lives in the future.”*
>
> *(Bolivia student, translated from Spanish)*
>
> *“Seeing how drugs act at a microscopic level in cells.”*
>
> *(Bolivia student, translated from Spanish)*

These comments suggest that the IoT-enabled imaging technology is sufficient to expose students to complex experiments while fostering an enjoyable and interesting experience and that the Picroscope imaging viewer is easy to use for high school students. These responses also reveal that the relevance of the scientific topic is an important factor for generating interest in the experiment and should be considered for future projects.

## 4.4. Remote PBL Is as Effective as In-Person PBL at Increasing Interest In Science

In addition to the STEM identity questions, at the end of the program, we gave our Bolivian student participants a series of questions related to their attitudes towards STEM and our remote PBL program. The questions had all previously been used to assess enthusiasm for STEM after an in-person biology PBL course with another set of Bolivians [1], and thus can be directly compared between the two cohorts.

We first assessed their attitudes towards science with the survey included in Supplemental Table 1. On questions relating to overall enthusiasm for science and interest in science careers, students all responded positively. The one question where the group answers differed was “hard work will help me be successful in science”, where 67% of the remote Bolivian cohort responded “strongly agree” compared to 90% in the in-person cohort (Figure 8A). On questions regarding feelings toward the program they participated in, all participants responded similarly, reporting positive feelings towards these programs (Figure 8A).

**Figure 8:**
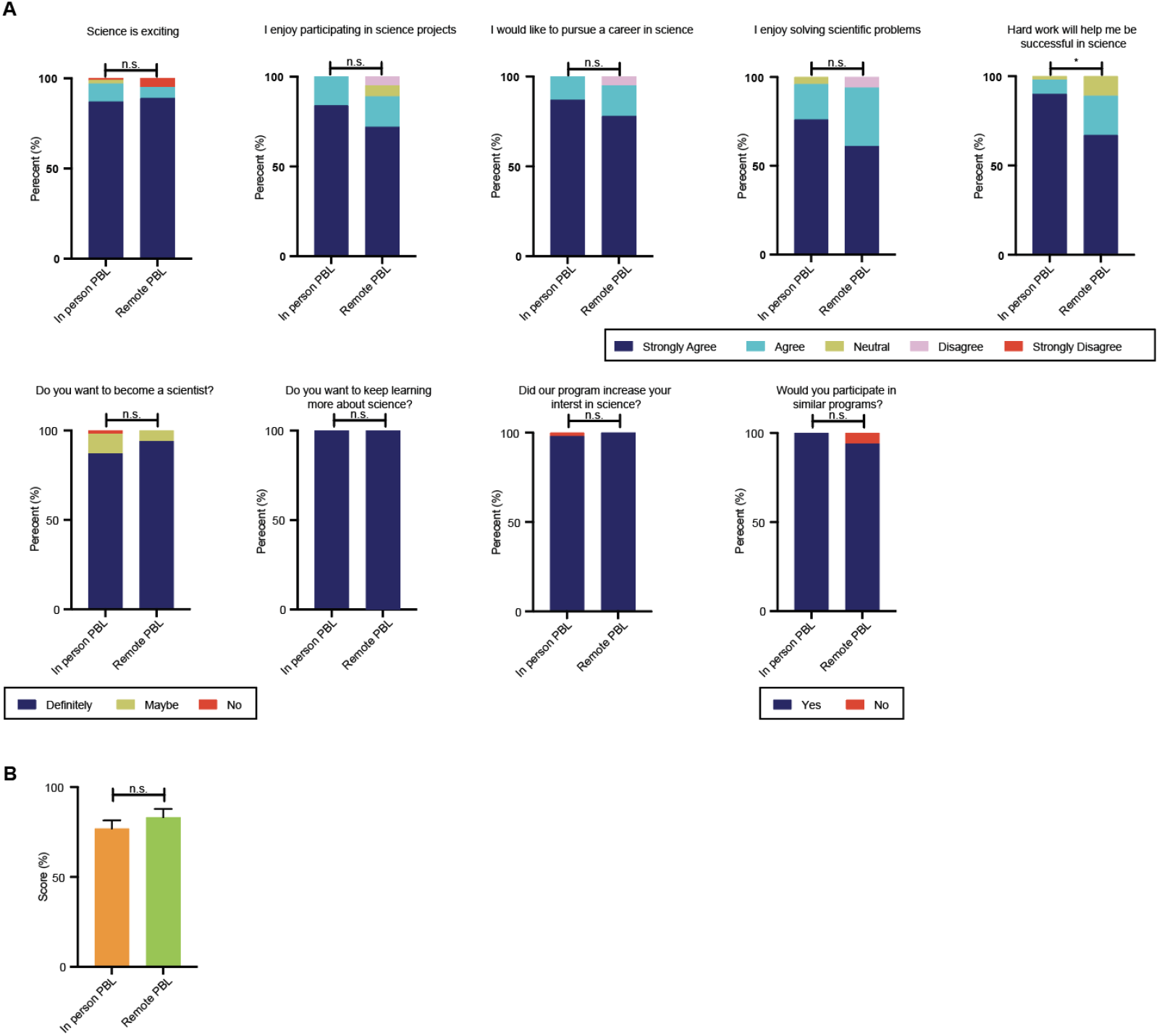
Remote PBL is as effective as in-person PBL at increasing interest in science and understanding of the scientific method. (A) Student’s levels of agreements used to assess enthusiasm for STEM after in-person and remote PBL courses. In person PBL is from Ferreira et al., 2019 [1]. Cohort sizes: In person PBL n = 92, Remote PBL n = 18. Mann Whitney test. * = p <0.5, n.s. = not significant. (B) Mean score received on a test related to the scientific method, administered following the remote or in-person PBL course at the Catholic University of Bolivia. Cohort sizes: In Person PBL n = 24, Remote PBL n = 17. Student’s t-test. n.s. = not significant. Error bars represent SEM. (D).

These results show that among students with similar attitudes towards science, remote PBL is as effective as in-person PBL at increasing interest in science and that the students enjoyed participating in the remote project.

## 4.5. Remote PBL Leads to Comparable Understanding of the Scientific Method to In-Person Instruction

To address whether our remote PBL approach leads to similar learning outcomes as in-person instruction, we tested our group in Bolivia with a fully in-person section of the same course. The students were tested on six questions related to the scientific method, including experimental design, data analysis, and outlier identification (Supplemental Note 6). We found no statistically significant differences between the group that had undergone remote PBL when compared to the in-person instruction (Figure 8B, in person = 76.96 +/−22.35%, n = 24; remote= 83.17 +/−19.45%, n = 17; p = 0.36), suggesting that remote experimentation is as effective at teaching the scientific method compared to in-person instruction.

## 5. Discussion

Despite significant investments in education, Latinx people continue to be underrepresented in the sciences. Education technology ‘megaprojects’ such as the One Laptop per Child project and EdX have shown that access to technology alone is ineffective at motivating underrepresented students [61, 62]. Therefore, novel approaches that can integrate scalable technologies with proven successful teaching methodologies are needed to target students effectively. Moreover, technologies that could effectively be deployed at scale should, in principle, be adaptable to various contexts, curricula, and languages [63]. Our framework uses IoT-enabled microscopy to complement high school and college biology courses for student populations in the United States, Bolivia, Brazil, Spain, Colombia, and Mexico. This framework allowed the instruction of the students in their native language using PBL and was adapted to a variety of real-world projects, such as caffeine consumption in students, agriculture chemical exposure, and pharmacology.

Currently, most studies in education in underrepresented groups are based on relatively small geographical locations. For example, studies in Latinx populations tend to be based on immigrants or immigrant-descent students living in the United States [64, 65]. Results are often extrapolated to Latinx populations worldwide, assuming that the populations living in the United States are representative of the native populations. Yet, previous work from our group has shown that, at least for biology education, results of studies in Latin America differed significantly from expected based on literature [1]. However, direct comparative studies between groups are difficult due to logistical and systematic difficulties, such as language barriers, differences in teacher training, and variations in academic calendars. Here, we took advantage of IoT-enabled microscopy to compare the effects of remote PBL between groups. For example, we found that Bolivia students had stronger STEM identities than their California counterparts (Figure 7). As strong STEM identity is a predictor of future STEM career choice [66], understanding the roots of these differences is important to better target education approaches that can increase diversity in STEM.

A unique aspect of this remote PBL paradigm is that it allows students to run experiments using materials that would otherwise be inaccessible due to their hazardous potential or the difficulty of transporting them to remote locations. For example, the AP Biology and college general biology curricula include content on mammalian cellular biology [67]. Yet, experiments in those courses have been limited in scope, focusing on microbiology or plant biology [68]. Students are usually not exposed to experimental mammalian cell and tissue culture until upper division college courses [69]. Given the differences in cellular biology between life kingdoms, there is a disconnect between theory and practice in high school and lower-level college biology courses. We addressed this disconnect by creating a context-informed project centered on culture of neuroblastoma cells and the effects of various drugs on those cells. These “clinical trials in a dish” expose students to many facets of science work and introduce them to career paths in STEM that may have been unknown to them, including biochemistry, analytical chemistry, and bioinformatics. We expect that this will be important for increasing diversity in STEM. Most surveyed students self-reported that this program positively impacted their interest in science and pursuing scientific careers.

Our future vision for the PBL remote experimentation framework is to create a community environment with massive open online experimentation. Many science labs and classrooms can collectively enroll in a session to connect students worldwide while still having them benefit from learning in a small classroom. Each session would focus on a specific experimental question. All students would crowd-source ideas about the procedure, launch experimental conditions, share and analyze data, and present their findings in a culminating event. Stay informed on future updates at https://braingeneers.gi.ucsc.edu/.

On the technical side, a future direction would be to connect more sensors and controllers to each experiment, such as devices that measure electrical activity, automated drug delivery, and light stimulation. Another exciting future direction would be to improve the software visualizations. An immersive online environment could be created in 3D through AR/VR (Augmented Reality/ Virtual Reality), where students use their phones to manipulate the experimental equipment and models. The manipulations they perform in a VR environment could change experimental devices in the lab connected through IoT. We are eager to continue exploring the potential of this technology to facilitate ever-improving educational experiences for everyone.

## 6. Supplemental Materials

## ACKNOWLEDGMENTS

We would like to thank all the students who participated in the user studies and whose excitement inspired this work. In addition, we are thankful to Zia Isola for her support setting up the groundwork for this project. We would also like to thank Sofie Salama for her critical feedback on this manuscript. This work was supported by the Schmidt Futures Foundation SF 857 (D.H. and M.T.), the National Human Genome Research Institute under Award number 1RM1HG011543 (D.H. and M.T.), and the the National Science Foundation under award number NSF 2134955 (D.H., and M.T.). D.H. is an investigator with the Howard Hughes Medical Institute. The content is solely the responsibility of the authors and does not necessarily represent the official views of the Schmidt Futures Foundation, NIH, NSF or HHMI.

## AUTHOR CONTRIBUTIONS

P.V.B., R.S., A.K.W., K.V., V.T.L., R.H., M.A.T.E., R.W., S.T.M., F.A., P.V., L.E.A.-A. and M.A.M.-R. taught the courses. P.V.B., K.V., V.T.L., M.A.T.E., D.F.P., D.E. and Y.M.R. developed the technologies used for remote education. G.M. provided administrated support in the United States. N.M.D. and L.E.A.-A. provided administrative support in Bolivia. N.M.D. provided admnistrative support with Science Clubs International. A.B. and T.S. provided technical support. S.K., D.H., M.T., and M.A.M.-R. supervised the work. All authors contributed to the writing of the manuscript.

## COMPETING INTERESTS

P.V.B., K.V., V.T.L., Y.M.R., D.H. and M.T. have submitted patent applications related to Internet-of-Things-enabled microscopy. M.A.M.-R. is a cofounder of Paika, a company for remote people-to-people interactions. The authors declare no other conflicts of interests.

