## Supplemental Materials for "Cloud-controlled microscopy enables remote project-based biology education in Latinx communities in the United States and Latin America"

Supplemental Figure 1: Bolivian students' presentations
Supplemental Figure 2: Answers of student cohorts to the RIS-STEM instrument
Supplemental Figure 3: Different student cohorts report similar effects to IoT-enabled PBL
Supplemental Table 1: Student Survey Questions effects of small molecules on neuroblastoma cells.
Supplemental Note 1: Data in Biology Activity
Supplemental Note 2: Model Organisms Activity
Supplemental Note 3: Experimental Design Activity
Supplemental Note 4: Performing Experiments Activity
Supplemental Note 5: Science Communication Activity
Supplemental Note 6: Scientific method test
Supplemental Video 1: Remote Image Viewer Tutorial
Supplemental Video 2: Alisal High School students' presentations reporting their results of the

Available upon request to the corresponding author

**Supplemental Figure 1. Bolivian students' presentations.** (A) Bolivian students presented the results of their observations on the effects of small molecules on neuroblastoma cells in a local science fair.

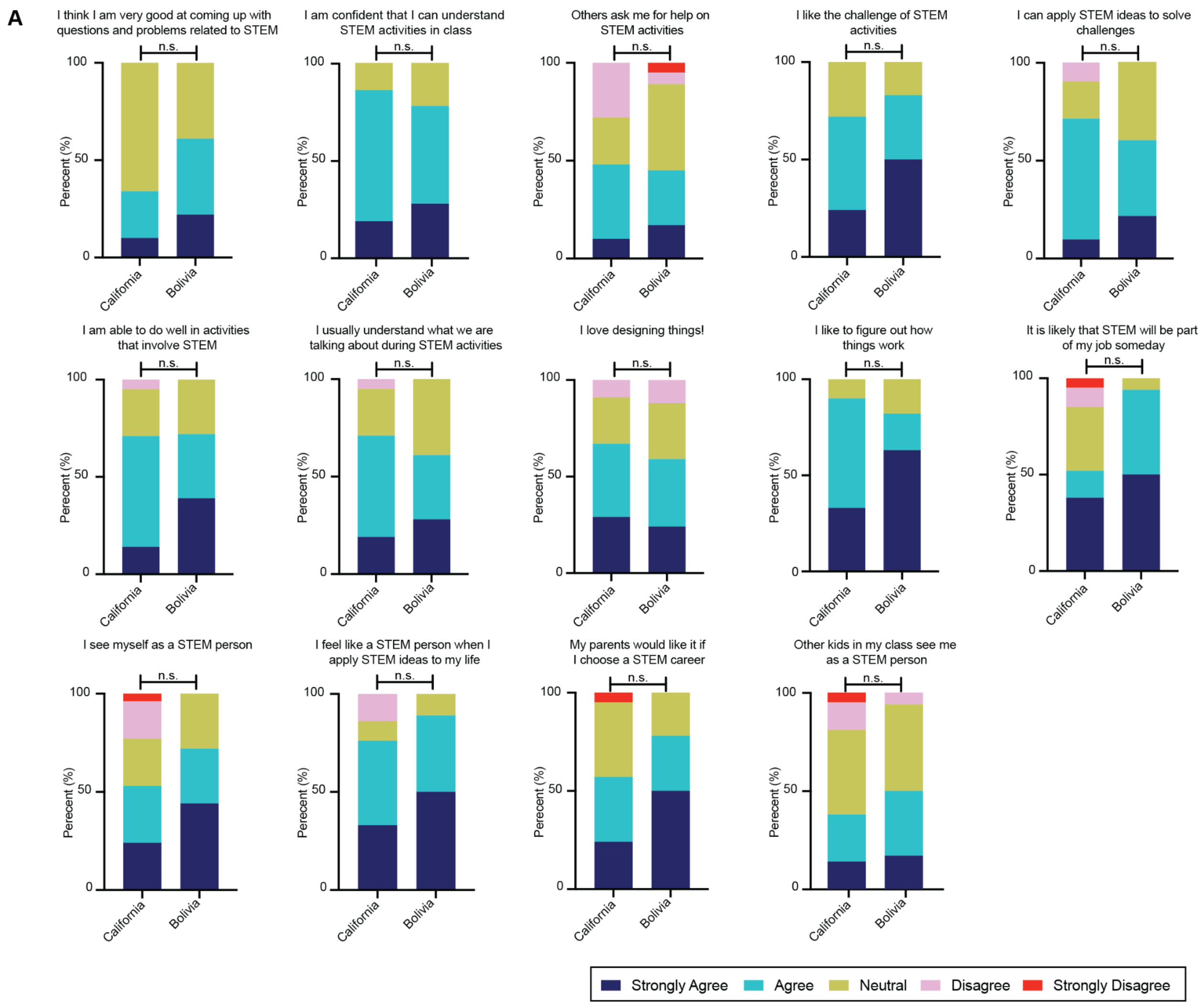

**Supplemental Figure 2. Answers of student cohorts to the RIS-STEM instrument.** (A) Remaining answers to the RIS-STEM instrument in complement to Figure 5. Cohort sizes: California n = 21, Bolivia n = 18. Mann Whitney test. n.s. = not significant.

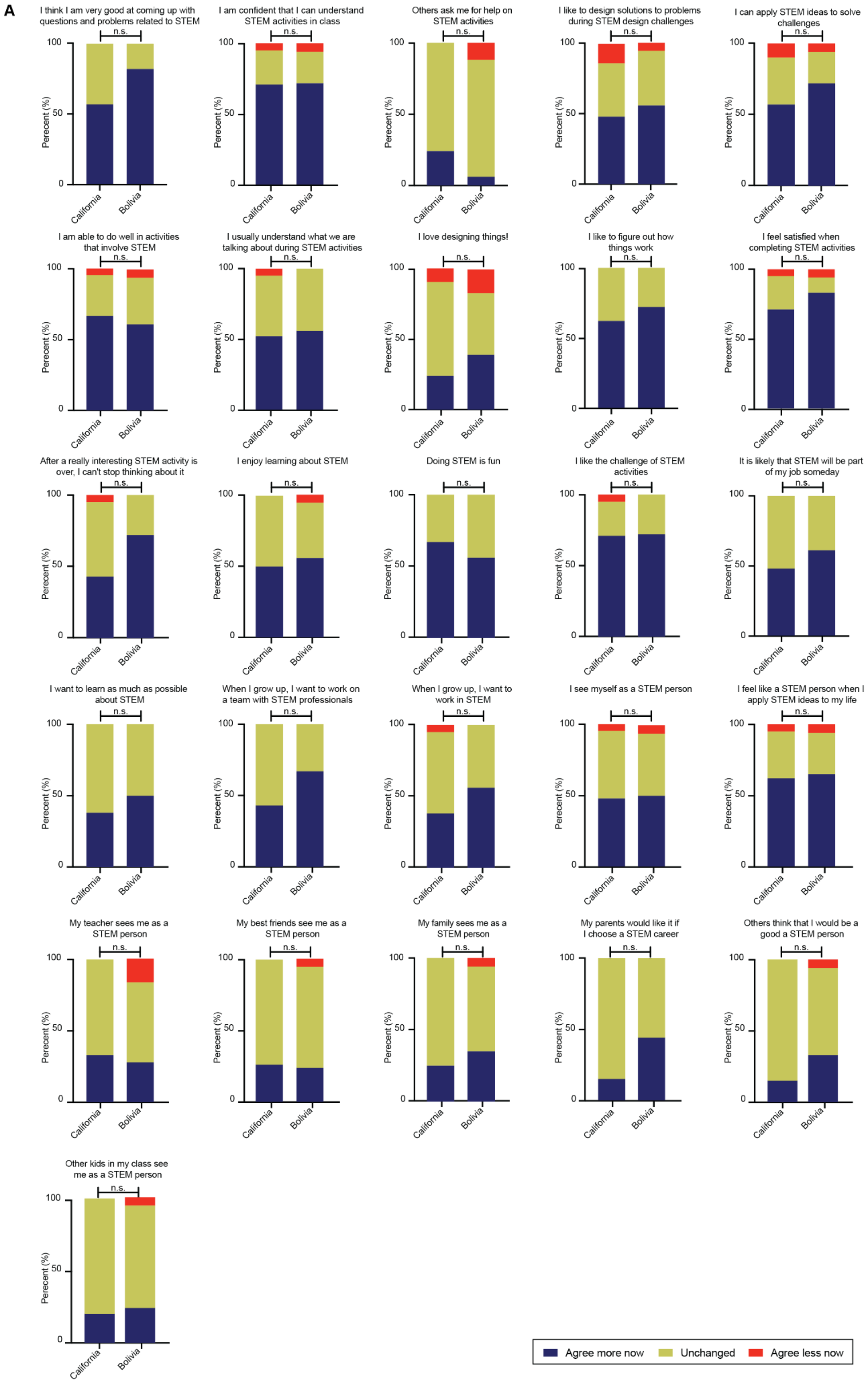

**Supplemental Figure 3. Different student cohorts report similar effects to IoT-enabled PBL.** (A) Students were asked how the program impacted their answers to the RIS-STEM instrument. Cohort sizes: California n = 21, Bolivia n = 18. Mann Whitney test. n.s. = not significant.

#### Supplemental Table 1: Student Survey Questions

| Free response | Attitudes towards STEM (Ferreira et al., 2019) | Attitudes towards STEM (Ferreira et al., 2019) | Attitudes towards our program | STEM professional overlap (McDonald et al., 2019) | STEM identity questions (Paul et al., 2020) |
|---|---|---|---|---|---|
| 1) What was the most interesting or useful thing you feel you learned? | Rate your level of agreement with the following statements: (Options are strongly agree, agree, neutral, disagree, strongly disagree) | Answer the following questions: (Options are no, maybe, definitely) | Answer the following questions: (Options are yes, no) | 1) Select the picture that best describes the current overlap of the image you have of yourself and your image of what a STEM professional is. (See Figure 6 for answer options) | Rate your level of agreement with the following statements: (See Figure 7 and Supplmental Figure 1 for questions, or refer to Paul et al., 2020. Answer options are strongly agree, agree, neutral, disagree, strongly disagree) |
| 2) How user friendly was the imaging viewer (webpage in order to access the images) for you? Do you have any suggestions/changes about how to improve it? | 1) Science is exciting. | 1) Do you want to become a scientist? | 1) Did our program increase your interest in science? | 2) How do you think participating in this project influenced your answer to the question above on your image of a STEM professional? (Options are "My self-image overlaps MORE, THE SAME, or LESS with STEM Professional than it did before") | Now, consider how you would have answered question 6 BEFORE participating in this project. Did your participation in this project change your answers to the statements in the above questions? Select how each changed after participating. (Options are agree more now, agree less now, unchanged) |
| 3) Did you enjoy being able to watch a live experiment in almost real time? If yes, what did you like about it? | 2) I enjoy participating in science projects. | 2) Do you think you can succeed in science? | 2) Would you participate in similar programs? | | |
| | 3) I would like to pursue a career in science. | | 3) Do you want to keep learning more about science? | | |
| | 4) I enjoy solving scientific problems. | | | | |
| | 5) Hard work will help me be successful in science. | | | | |

### How Do I Interpret My Scientific Data?

#### Learning Goal: Understanding the data and the secrets behind it

##### <u>Day 1: Types of Data and Intro to Programming</u>

**1. Quantitative:** Data in the form of counts or numbers.

**Examples:** Height, weight, time, size, temperature

**2. Qualitative:** Data that is non-numerical and collected through methods of observation.

**Examples:** Appearance, scent, feelings, textures

##### <u>How do I analyze my data?</u>

**Quantitative data:**
- Compare statistics (examples: mean, median, mode)

**Qualitative data:**
- Grouping by categories
- Hint: make your qualitative data into quantitative data!

**Graphs & Charts**

**Line graph:** A function that changes depending on the relationship between dependent and independent variables. Example:

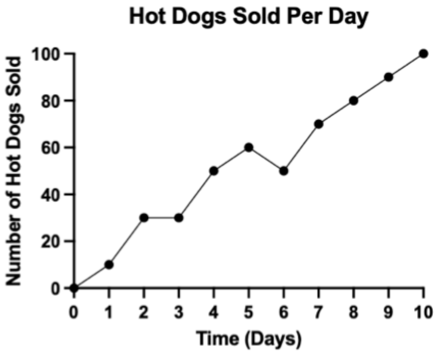

**Scatter plot:** Explores the association between two variables, which could be quantitative or qualitative. Example:

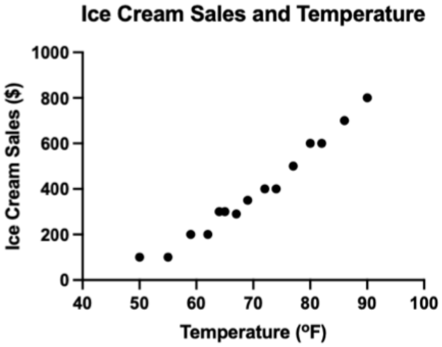

**Bar graph:** Shows quantitative differences between categories. Normally used to quantify categorical data. Example:

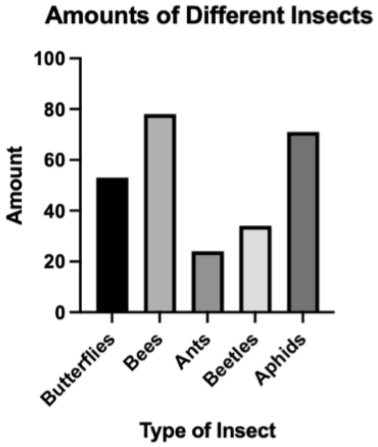

**Histogram:** Shows the frequency of data divided into groups or “bins”. Example: (here, the bins are are the heights of the trees)

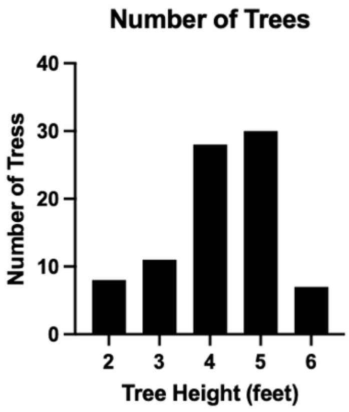

**Box-and-whisker plot:** Shows variation within your data. Example:

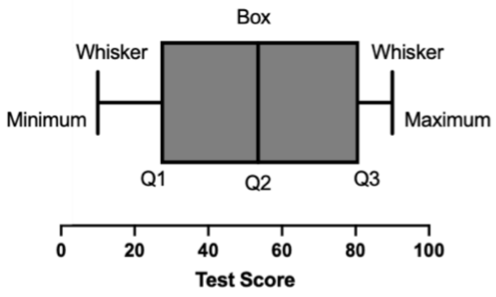

Five important parameters of a box-and whisker plot:

1. Maximum
2. Upper Quartile (Q3, 75%)
3. Median (Q2)
4. Lower Quartile (Q1, 25%)
5. Minimum

**Pie chart:** Shows different parts of the whole, each slice represents the proportion of one category. Example:

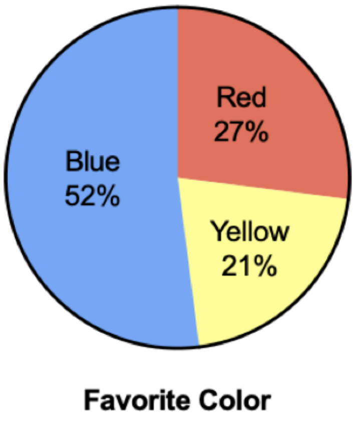

**When selecting the type of graph for your data, think about:**

1. **What type of data you have:** Is it categorical, quantitative?
2. **Purpose of the data:** What do you want to show/compare/interpret?

**Now that we know about the different types of graphs, let's make some ourselves!**

#### Intro to programming and data visualization

Instructions: Get in break-out rooms of groups of 2-3 students.
Take 30 mins to work with your group on the activities. Work collectively on the notebook and be ready to share your answers with your classmates!

**Colab notebook:**
https://colab.research.google.com/drive/1UG_lHT3jQ-cC2fGCYbbnDyCs_Q1TC5xK?usp=sharing

#### Day 2: Generating Data

Note: if you do want to do the actual experiment, the sample data on the next page can be used to go straight to the data visualization activity (Day 3).

---

Learn-Biology.com

### Gummy Bears, Raisins and Osmosis

**1. Setup (Day 1):** Label a piece of masking tape with your name. Use that tape to tape a gummy bear and a raising to the outside of a plastic cup. Place a second gummy bear and a second raisin into the cup. Fill the cup about ½ way with water and let sit overnight.

**Day 2. Data collection and analysis**

**A. Visual Observations**: How do the gummy bears and raisins look different?

______________________________________________________________________________

______________________________________________________________________________

______________________________________________________________________________

**B. Numerical Data. Carefully pour off all water before weighing. Zero the scale each time.**

| | A | B | C | D |
|---|---|---|---|---|
| **Note: you'll have to work with another group to get one data point below** | **Individual data (mass in grams)** | **Total class data for gummies (mass in grams)** | **# of gummies** | **Average class data (Total mass/# gummies )** |
| **Dry Gummy Bear** | | | | |
| **Gummy Bear in tap water** | | | | |
| **DRY RAISIN** | | | | |
| **RAISIN IN TAP WATER** | | | | |

***Graph: Class data for our gummy bear lab***

| **MMASS in grams** | | **Gummy Bear** | | | | **RAISIN** | |
|---|---|---|---|---|---|---|---|
| | | **DRY** | | **Tap water** | | **DRY** | **TAP** |

**C. Using the concept of *diffusion*, can you explain what happened?**

_____________________________________________

_____________________________________________

_____________________________________________

_____________________________________________

_____________________________________________

_____________________________________________

_____________________________________________

_____________________________________________

_____________________________________________

_____________________________________________

**Sample Data**

| 6 bears per sample | | | |
|---|---|---|---|
| | Dry mass | 10% sucrose mass | Tap water mass |
| Group 1 Oscar | 3.08g | 12.14g | 8.90g |
| Group 2 Karizma | 3.07g | 11.47g | 9.78g |
| Group 3 Bri | 3.09g | 8.43g | 9.29g |
| Group 4 Naomi | 3.17g | 11.86g | 9.13g |
| Group 5 Emilie | 3.01 g | 11.52 g | 9.08 g |
| Group 6 Adrian | 3.07g | 11.48g | 8.77g |
| Average | 3.08 | 11.15 | 9.16 |

### Day 3: Data Interpretation

Today we are doing a hands-on group activity. We will be analyzing your data from the gummy bear experiment in different ways and then presenting them to your peers in an engaging way.

We've talked about the scientific method and learned about some tools to look at your data but now it's time to visualize your data and present it to your peers. This is where you get to show off your methods and results and explain your data to someone who has never seen your project before.

It's important to take a step back from what you did to try to see the project as someone who might not know about it. Explain your hypothesis, and methods, and visualize your data in a simple way so that anyone could understand what you did, how you did it, and what the results were. Make sure to highlight any cool results, outliers, and where your data followed and fell short of your hypothesis. The more clearly you present your experiment and data, the better someone will be able to use it to build on your research.

#### 1. Let's graph!

Visualize and present your data from the gummy bear experiment in a distinct way. Recall the examples from before.

- This is where you get to show off your results and explain your data to someone who has never seen your project before
- Practice making graphs
- Decide what type of graph will have the most impact
- Make a 1-minute video to teach your peers about how you represented your gummy bear data using a graph. See step 2 for guidelines:

#### 2. Make a Presentation Video

Tik-Tok or Instagram format:
20 sec: Show your graph - how you labeled it and what it describes.
10 sec each: Choose 3 types of graphs and explain why they are useful (it's best for what kind of experiment/data?)
10 sec: Why is data visualization important for scientists?

#### 3. Contest: vote on which team had the best video!

**Supplemental Note 2: Model Organisms Activity**

### Group practice questions

**Question 1:** You hypothesize that lead exposure in drinking water causes a decrease in fertility.

Design an experiment to test your hypothesis using one of the model organisms discussed today. Explain why you chose that specific model organism.

### Group practice questions

**Question 2:** You discover a new protein with a similar sequence and structure to a protein called SALL1. SALL1 is necessary for limb development of the human embryo. You hypothesize that the function of your new protein is identical to SALL1.

Design an experiment to test your hypothesis using one of the model organisms discussed today. Explain why you chose that specific model organism.

### Group practice questions

**Question 3:** You discover a new gene that you name Alisal. You hypothesize that Alisal is essential for neuronal development and cognitive ability.

Design an experiment to test your hypothesis using one of the model organisms discussed today. Explain why you chose that specific model organism.

**Supplemental Note 3: Experimental Design Activity**

### Question

How do certain drugs affect cancer cells?

### Background

**What do these drugs do?**

Retinoic Acid:

Neurodazine:

Primocin:

### Conditions

Class splits up into 3 groups (retinoic acid, neurodazine, primocin)

Each group will form a hypothesis:
"If ... (independent variable)... then ...(dependent variable).. because ...(rationale)..

Things to think about:

- Do the cells divide?
- Do the cells move? Stop moving?
- Do the cells change shape? Do they become neurons?
- Do the cells die?
- How can this all be measured?

#### Retinoic Acid Group

**Hypothesis**:

#### Neurodazine Group

**Hypothesis**:

#### Primocin Group

**Hypothesis**:

**What will you look for in the data and how will it relate to the hypothesis? (hint: what defines cancerous cells?)**

### Variables

What data are we measuring?

What are your variables? (independent variable, dependent variable, controls)

### Materials and Methods

#### Materials:

- Microscope
- Cancer cells
- 24-well plate
- Media to grow the cells in
- Drug to test

#### Well plate layout:

Use this as a template to indicate where each condition will go
Include at least 3 replicates of the drug and control treatments

### Methods for the experiment:

1. Put cells into the wells
2. Add drugs to cells, make sure the same amount is given to each well
3. Take pictures of the cells every hour through the duration of the experiment (3 days)

### Methods for the observation/data collection:

1. Obtain picroscope images from web interface (control and drug treated cells)
2. Observe how the cells react to the drugs each day
3. Record your observations

**Supplemental Note 4: Performing Experiments Activity**

### Data

#### Group 1

**Retinoic: Well # ____**

| | **Cell identification** (How do the cells look? Are they cancerous cells or neurons?) | **Cell Size compared to control** (qualitative) | **Total cell count** (quantitative, number) | **Number of neurons vs cancerous cells** (quantitative, proportion) |
|---|---|---|---|---|
| **Day 1** | | | | |
| **Day 2** | | | | |
| **Day 3** | | | | |

**Control: Well # ____**

| | **Cell identification** (How do the cells look? Are they cancerous cells or neurons?) | **Total cell count** (quantitative, number) | **Number of neurons vs cancerous cells** (quantitative, proportion) |
|---|---|---|---|
| **Day 1** | | | |
| **Day 2** | | | |
| **Day 3** | | | |

#### Group 2

**Neurodazine: Well # ______**

| | **Cell identification** (How do the cells look? Are they cancerous cells or neurons?) | **Cell Size compared to control** (qualitative) | **Total cell count** (quantitative, number) | **Number of neurons vs cancerous cells** (quantitative, proportion) |
|---|---|---|---|---|
| **Day 1** | | | | |
| **Day 2** | | | | |
| **Day 3** | | | | |

**Control: Well # ______**

| | **Cell identification** (How do the cells look? Are they cancerous cells or neurons?) | **Total cell count** (quantitative, number) | **Number of neurons vs cancerous cells** (quantitative, proportion) |
|---|---|---|---|
| **Day 1** | | | |
| **Day 2** | | | |
| **Day 3** | | | |

#### Group 3

**Panobinostat/Primocin: Well # ______**

| | **Cell identification** (How do the cells look? Are they cancerous cells or neurons?) | **Cell Size compared to control** (qualitative) | **Total cell count** (quantitative, number) | **Number of neurons vs cancerous cells** (quantitative, proportion) |
|---|---|---|---|---|
| **Day 1** | | | | |
| **Day 2** | | | | |
| **Day 3** | | | | |

**Control: Well # ______**

| | **Cell identification** (How do the cells look? Are they cancerous cells or neurons?) | **Total cell count** (quantitative, number) | **Number of neurons vs cancerous cells** (quantitative, proportion) |
|---|---|---|---|
| **Day 1** | | | |
| **Day 2** | | | |
| **Day 3** | | | |

#### Supplemental Note 5: Science Communication Activity

### Scientific Presentation Outline

**Background:**

1. Introduce what you studied.
    a. For example, if you were researching a drug, describe how the drug is used in patients and how it works.
2. State the knowledge gap. In other words, describe what is not known about the thing you studied.
    a. For example, maybe it is not known how the drug you studied affects cancer cells, so you want to test this.
3. Tell the audience why you studied this and why it is important!

**Hypothesis:**

1. State your hypothesis.

**Methods:**

1. Describe the steps taken to test your hypothesis.

**Results:**

1. Present your results using pictures, graphs, words, etc.
    a. Hint: the fewer written words on your results slides, the better. A picture is worth a thousand words!

**Conclusions:**

1. State what you concluded from the results.
2. Describe how this conclusion could be applied to real life situations.
    a. For example, if you found that your drug slowed cancer cell growth, maybe this drug could be used in cancer patients in the future.

**Supplemental Note 6: Scientific method test**
(Translated from Spanish)

**Question 1: The graph below shows the effect of temperature on the population size of three different species of fish.**

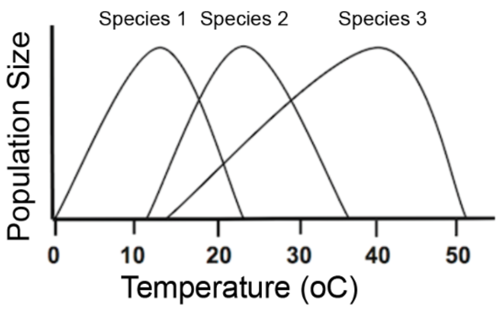

**Which species would thrive at a temperature of 40ºC?**

a) Species 3
b) Species 2
c) None of the species can tolerate this temperature
d) Species 1

**Question 2: An ecologist calculated the average annual rainfall in a local valley. His data is in the following table:**

| Month | Precipitation (mm) |
|---|---|
| January | 16 |
| February | 20 |
| March | 29 |
| April | 23 |
| May | 15 |
| June | 7 |
| July | 3 |
| August | 1 |
| September | 2 |
| October | 4 |
| November | 6 |
| December | 11 |

**In which month was there more rain?**

a) November
b) April
c) August
d) March

**Question 3: A biology student wanted to determine if there is a relationship between resting heart rate and height. She gathered information from 12 classmates and constructed the following graph.**

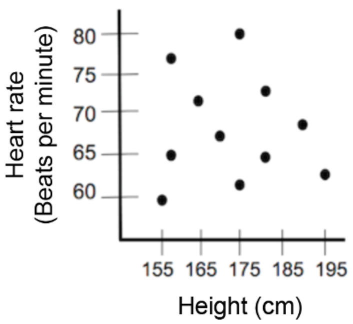

**Based on the data, which of the following is true?**

a) As height increases, resting heart rate decreases.
b) As height decreases, resting heart rate increases.
c) Height and resting heart rate are not directly correlated.
d) As height increases, resting heart rate also increases.

**Question 4: A biology student wanted to determine if there is a relationship between resting heart rate and height. She gathered information from 12 classmates and made the following table.**

| Height of the student (cm) | Resting heart rate (beats per minute) |
|---|---|
| 155 | 60 |
| 156 | 65 |
| 156 | 78 |
| 165 | 72 |
| 170 | 67 |
| 175 | 62 |

| | |
|---|---|
| 175 | 80 |
| 180 | 64 |
| 180 | 73 |
| 190 | 68 |
| 194 | 78 |
| 195 | 63 |

**Which of the following statements is best supported by the data in the table?**

a) The taller the student, the higher the resting heart rate.
b) Higher resting heart rate results in growth in students.
c) There is no direct correlation between height and resting heart rate.
d) The shorter the student, the higher the resting heart rate.

**Question 5: A class of biology students conducted an experiment on ants. Each group received 20 random ants. Each ant was placed the same distance from a dark-colored path that led to food and a bright-colored path that led to food. Five of the groups found that the ants chose the colored path more often. Three groups found that ants prefer the dark path, and two groups did not find that their ants had a preference. A student who proposed the hypothesis that ants are attracted to bright colors found that this hypothesis was supported by data from his group. How does the student's conclusion exemplify scientific bias?**

a) The group of students selected the ants that were most likely to support their hypothesis.
b) The students were separated into several groups to carry out the experiment, which skewed the results.
c) The sample size in the experiment was too small to produce valid results.
d) The student did not give importance to the data that did not support the hypothesis.

**Question 6: A biology student wanted to study the effect of a fertilizer on the growth of tomato plants. He placed four tomato plants of the same type in separate containers, each containing the same amount of soil. Each pot was watered with a different solution containing different amounts of fertilizer. The following table shows the average height of the plants at the end of a week. Plant group, amount of fertilizer (g), average height of plants (cm)**

| Plant group | Amount of fertilizer (g) | Average height of plants (cm) |
|---|---|---|
| 1 | 0 | 3.0 |
| 2 | 5 | 3.2 |

| | | |
|---|---|---|
| 3 | 10 | 4.4 |
| 4 | 20 | 5.0 |

**Which group is the control group?**

a) Group 2
b) Group 4
c) Group 3
d) Group 1

**Supplemental Video 1: Remote Image Viewer Tutorial**

Available upon request to the corresponding author

**Supplemental Video 2: Alisal High School students' presentations reporting their results of the**

Available upon request to the corresponding author
